# Influence of light on particulate organic matter utilization by attached and free-living marine bacteria

**DOI:** 10.1101/537415

**Authors:** Laura Gomez-Consarnau, David M. Needham, Peter K. Weber, Jed A. Fuhrman, Xavier Mayali

## Abstract

While the impact of light on primary productivity in aquatic systems has been studied for decades, the role light plays in the degradation of photosynthetically-produced biomass is less well understood. We investigated the patterns of light-induced particle breakdown and bacterial assimilation of detrital C and N using ^13^C and ^15^N labeled freeze-thawed diatom cells incubated in laboratory microcosms with a marine microbial community freshly-collected from the Pacific Ocean. Particles incubated in the dark resulted in increased bacterial counts and dissolved organic carbon concentrations compared to those incubated in the light. Light also influenced the attached and free-living microbial community structure as detected by 16S rRNA gene amplicon sequencing. For example, bacterial taxa from the Sphingobacteriia were enriched on dark-incubated particles and taxa from the family Flavobacteriaceae and the genus Pseudoalteromonas were numerically enriched on particles in the light. Isotope incorporation analysis by phylogenetic microarray and NanoSIMS (a method called Chip-SIP) identified free-living and attached microbial taxa able to incorporate N and C from the particles. Some taxa, including members of the Flavobacteriaceae and Cryomorphaceae, exhibited increased isotope incorporation in the light, suggesting the use of photoheterotrophic metabolisms. In contrast, some members of Oceanospirillales and Rhodospirillales showed decreased isotope incorporation in the light, suggesting that their heterotrophic metabolism, particularly when occurring on particles, might increase at night or may be inhibited by sunlight. These results show that light influences particle degradation and C and N incorporation by attached bacteria, suggesting that the transfer between particulate and free-living phases are likely affected by external factors that change with the light regime, such as time of day, depth and season.

## Introduction

Photoheterotrophy is a major biogeochemical process in surface waters. Although this dogma is now recognized across the aquatic microbiology field (Béjà et al. 2000; Kolber et a. 2000), until the year 2000 all marine heterotrophic bacteria were classified as strict organic matter decomposers. Two major light energy transducing mechanisms are now identified in surface bacterioplankton: proteorhodopsin (PR) photoheterotrophy and aerobic anoxygenic phototrophy (AAP). PRs are the most abundant and widespread of the two. Notably, they are present and highly expressed in many bacteria including those from the SAR11 clade (Giovannoni et al. 2005, Gifford et al. 2013, Ottessen 2013, 2014) and most studies suggest that they are ubiquitous in nutrient-depleted waters (e.g. the Eastern Mediterranean Sea; Gomez-Consarnau et al., 2017, Dubinsky et al., 2017). However, PRs have also been identified in bacteria adapted to grow on organic matter rich particles such as the Flavobacteriia (Gómez-Consarnau et al. 2007, Gonzalez et al. 2008, Riedel et al. 2013, Maresca et al. 2018). These nutrient-rich particle-associated microenvironments also seem to be preferred by some AAP bacteria, such as the Rhodobacteraceae commonly found after seasonal phytoplankton blooms (e.g. Allers et al. 2007) and during outdoor cultivation of microalgae (Geng et al. 2016). One unanswered question is whether light energy from PR and AAP photoheterotrophy can be used by those microorganisms to optimize their resource utilization for the degradation of complex particulate organic matter.

Previous studies carried out in free-living communities strongly suggest that photoheterotrophy plays a role in dissolved organic matter (DOM) degradation. A study in the North Pacific subtropical gyre at station ALOHA reported bacterial production rates 48-92% higher in the light compared to the dark using radiolabeled leucine incorporation (Church et al. 2004). While the microbial groups responsible for the observed light-enhanced uptake rates were not identified, the authors attributed this observation to possible Synechococcus amino acid uptake. Similarly, Michelou et al. (2007) showed that light stimulated both amino acid assimilation and bacterial production; in this case the effect did not correlate with cyanobacterial abundances nor with the growth of photoheterotrophic AAP. Other field studies have reported that both Prochlorococcus and low nucleic acid (LNA) bacteria (often dominated by the PR-containing SAR11 bacteria clade) show a significant increase in amino acids uptake in the light compared to the dark (Mary et al. 2008, Gómez-Pereira 2013, Evans et al. 2015). These data indicate that photoheterotrophy can support dissolved organic matter processing. However, whether or not light can also stimulate the uptake or degradation of particulate organic matter is currently unknown.

Although no studies have thus far examined the role of photoheterotrophy in the degradation of algal-derived particles, there is ample literature about the role of particle degradation in ocean carbon cycling (reviewed by Simon et al. 2002). These studies have determined that particle-attached microbes are i) phylogenetically diverse, often (but not always) dominated by Bacteroidetes known to degrade polysaccharides, ii) exhibit high solubilization rates and iii) may release labile substrates into the surrounding water (Azam & Long 2001). Understanding the interplay between free-living and particle-attached bacteria is key to further constrain the role of particles in elemental cycling. This question has been explored previously in dark incubations of live diatoms in laboratory rolling tanks (Passow et al. 2003), showing initial increases of biomass in particulate aggregates, further suggesting that the attachment of initially free-living bacteria may bring carbon from the free-living to the particulate phase. Bacteria have also been shown to rapidly attach and detach from particles but become irreversibly attached after some period of time (Kiørboe et al. 2003). Particle-attached bacteria exhibit higher bacterial protein production and higher protease activity per cell than when free-living (Grossart et al. 2007). In one example, polysaccharide-degrading bacterial strains could grow on laminarin (one of the most common carbohydrates in the sea) and released glucose and larger glucans in this process (Alderkamp et al. 2007). Yet, no previous reports have documented POM being released and subsequently incorporated by free-living bacteria; furthermore, the role of light and photoheterotrophy in the process of particulate breakdown remains uncharacterized.

In the current study, we address some of these unanswered questions using laboratory incubations of a microbial community that started as primarily free-living in the presence of stable isotope labeled phytodetrital particles. We aimed to test the hypothesis that photoheterotrophy enhances the transport of organic compounds from particles into bacterial cells, which would be consistent with light promoting the degradation of particulate organic matter (POM). In particular, this process is expected to be more important in taxa known to be particle-attached photoheterotrophs such as members of Flavobacteria and Rhodobacterales. We carried out laboratory incubation experiments using freshly collected free-living marine bacterial communities with diatom-derived particles to link the activity of both attached and free-living bacteria to the incorporation of particulate material. Dead, isotopically labeled phytoplankton cells were mixed with marine microbial communities in rolling bottles to increase cell collisions creating larger particles while also keeping them constantly suspended (Shanks & Edmondson 1989). We quantified incorporation of this particulate material by individual bacterial taxa using Chip-SIP, a method that uses stable isotope labeling, hybridization of RNA to phylogenetic microarrays, and isotopic imaging of the arrays using high-resolution secondary ion mass spectrometry (SIMS) with a Cameca NanoSIMS 50 (Mayali et al. 2012).

## Materials and Methods

### Production of ^15^N and ^13^C labeled diatom-derived particles

We used ^15^N and ^13^C labeled phytodetrital particles from the diatom Thalassiossira pseudonana CCMP 1335 as a representative of phytoplankton-derived organic matter that is likely to reflect the composition of natural particles in the ocean, and dead algal detritus in algal biofuel production ponds. Cultures of the Thalassiossira pseudonana CCMP 1335 were grown in F/2 medium (Guillard 1975) with 15NO3-under continuous light at 20° C. To maximize the incorporation of labeled carbon, bottles containing the medium were tightly shut immediately after autoclaving to prevent equilibration with atmospheric air, and once cooled were bubbled with N_2_ gas for 10 minutes to remove ^12^CO_2_ . 99% atm% ^13^C labeled sodium bicarbonate (NaH^13^CO_3_) was added at a concentration of 2 mM. A 100 μL inoculum of T. pseudonana already growing in ^13^CO_2_ and ^15^NO_3_ medium was added and the cultures were incubated with the bottle cap shut. After early stationary phase of growth was achieved after approximately two weeks, cultures were centrifuged at 1200 g, the liquid removed, and the pellet washed twice in filtered seawater, and frozen at −80 °C. Pellets were thawed and refrozen four times and washed in filtered artificial seawater to remove DOM from lysed cells, and frozen again before the start of the incubation experiment.

### Experimental setup, DOC analyses and bacterial counts

Surface seawater from the Coastal Pacific was collected at the San Pedro Ocean Time Series SPOT Microbial Observatory site (33° 33’N, 118° 24’W) on May 22^nd^ of 2013 and taken to the laboratory within four hours in a temperature controlled container. The seawater was subsequently gently filtered to remove grazers and 5 *μ*m particles (5 μm pore-size polycarbonate filters, Millipore corp., Billerica, MA), and then distributed into 10 polycarbonate 2-liter cylindrical bottles (Nalgene). SIP-addition treatments were enriched with 0.19 g/L dry weight diatom-derived ^13^C and ^15^N labeled particles and incubated in the light (at 150 μmol m-2 s-1) or in the dark in triplicate at 18°C in a rolling bottle setup at 0.5 rpm (CellRoll, Inc). Duplicate control treatments with no added particles were incubated in the same temperature controlled chamber in the light or dark in static bottles. Samples for cell counts, dissolved organic carbon (DOC), DNA and RNA were collected after 72 hours in addition to T0. All material used was previously acid washed with 1% HCl and rinsed in double distilled water.

Samples for DOC were collected by filtering 100 ml of sample through 0.2 *μ*m pore-size polycarbonate filters (47 mm, Millipore corp., Billerica, MA). All material in contact for those samples was pre-rinsed with 1M HCl. DOC samples were sent to the Nutrient Analysis Lab at University of Maryland for analysis. Samples for bacterial counts were preserved in 10% formalin and kept at 4°C until processing. Two milliliters of the formalin-fixed samples were stained using acridine orange (Hobbie et al. 1977), filtered onto 0.2 *μ*m pore-size black polycarbonate Track-Etched (PCTE) filters (25 mm, Fisher Scientific) and counted with epifluorescence microscopy. Samples for DNA and RNA were collected on 5 *μ*m pore size polycarbonate filters (47 mm, Millipore corp., Billerica, MA), representing large particles, henceforth referred to as “particulate”, and the filtrate sequentially collected on 0.2 *μ*m pore size filters (47 mm diameter Durapore^®^ Membrane Filters, Millipore corp. Billerica, MA), representing free-living microbes and those attached to small particles, henceforth referred to as “free-living”.

### Nucleic acid extraction, and sequencing

RNA and DNA from particles (> 5 *μ*m) and free-living bacteria (0.2-5 *μ*m) from all replicates was extracted with the Qiagen AllPrep kit according to manufacturer’s instructions for bacterial nucleic acid, using 10 mg/mL lysozyme in the lysis step and including vortexing for 10 min. Bacterial diversity in the different treatments and size fractions was assessed by amplifying the V4 and V5 region of the 16S rRNA gene from the DNA extracts using primers 515F and 926R, and paired end 2 × 300 sequencing on MiSeq (Illumina)(Parada et al. 2015). Demultiplexed sequences were quality filtered with Trimmomatic trimming reads with the following settings: SLIDINGWINDOW:5:30, MINLEN:250. Paired end reads were then merged with usearch fastq_mergepairs with the following settings: -fastq_truncqual 5 -fastq_maxdiffs 0 -fastq_minmergelen 200 -fastq_minovlen 50. Primer sequences were removed with cutadapt. Chimeric sequences were identified in QIIME with identify_chimeric_seqs. py using a de novo and referenced based approach using the usearch61 method and the SILVA GOLD database. Operational taxonomical units (OTU) were determined using the uclust in QIIME (Edgar 2010) with 99% similarity cutoffs. The most abundant sequence within each OTU was selected as a representative sequence. Each representative sequence was assigned classification with QIIME assign_taxonomy.py against the SILVA 111 database. For identification of OTUs that were significantly more or less abundant in the different treatments and controls, we used the software packages phyloseq (McMurdie and Holmes, 2013) and DESeq2 (Love et al. 2014). Biom tables of read counts produced from the QIIME pipeline were imported into R via import_ biom of the phyloseq package. For samples where technical replicates were available (n=2), the reads from the replicates were summed. Then, the phyloseq object was converted to deseq2 objects via phyloseq_to_deseq2 for each statistical comparison of interest (light vs. dark, large vs. small, and control vs. experimental treatment). For determination of differential abundance between the experimental treatments (i.e., light vs. dark, and large vs. small size fraction), all of the different treatments of the other factor were used (that is, for example, data from both size fractions were used to determine significant shifts in light vs dark treatments). Significant differences were determined with the DESeq command using a “Wald” test and a “parametric” fitType (i.e., the default settings). Results were converted to a results table with the DESeq results command, retaining all results for further exploration (i.e., cooksCutoff=F). Significant shifts were identified as those with a Benjamini-Hochberg adjusted p-value < 0.05. Heatmaps were generated with the program “heatmap3” in R. Phylogenetic trees were generated by alignment with muscle (-maxiters 100, Edgar 2004) and maximum likelihood tree generation with phyml (Guindon et al. 2010). Those included amplicon representative sequences from the experimental taxa, together with relevant environmental sequences of previously reported phytoplankton bloom responders at SPOT (Needham et al. 2016) and whole genome sequenced bacteria.

### Probe design

We aimed to target organisms commonly found at SPOT as well as those that become enriched on diatom-derived particles. For the former, sequences collected at SPOT were obtained from published literature (Brown et al. 2005, Needham et al. 2013. For the latter, we used probes from a previous Chip-SIP study that targeted dead T. pseudonanaparticleattachedbacteria(Mayalietal. 2016). For probes targeting both diatom detritus and SPOT sequences, trimmed sequences were manually checked for quality and submitted to DECIPHER’s Find Chimeras web tool (Wright et al. 2012), aligned with the online SINA aligner (Pruesse et al. 2007) and imported into SILVA database SSU_NR ver. 103 in the ARB software (Ludwig et al. 2004), which included 263,000 small subunit ribosomal RNA sequences. Sequences were added to the global phylogeny and 148 unique operational taxonomic units (OTUs) were identified. For each OTU, a set of 25 probes targeting the OTU and closely related sequences were designed (Table S1) using the probe design function in ARB. In most cases, probes were allowed to perfectly match with less than 5 sequences outside the targeted group (preferably none, but this was not always possible). Also, in most cases we required that two base-pair changes on the probe sequence (i.e., mismatches) did not match more than 100 sequences outside the targeted group. Our approach was to have multiple probes for each taxon, including probes that target the same region but are offset by a few bases in either direction. This leads to a range of probe melting temperatures for each taxon, and decreases the likelihood that a single probe exhibits non-specific signal. The isotope enrichment of taxa was subsequently compared among one another in relation to the probe hybridization strength, as further explained below.

### Chip-SIP analyses

RNA samples were split: one fraction saved for fluorescent labeling, the other was unlabeled for NanoSIMS analysis. Fluorescent Alexafluor 532 labeling was carried out with the Ulysis kit (Invitrogen) for 10 min at 90°C (2 μL RNA, 10 μL labeling buffer, 2 μL Alexafluor reagent), followed by fragmentation. All RNA (fluorescently labeled or not) was fragmented using 1X fragmentation buffer (Affymetrix) for 10 min at 90°C before hybridization and concentrated by isopropanol precipitation to a final concentration of 500 ngμL-1. Glassslidescoatedwithindium tin oxide (ITO; Sigma) were coated with silane Super Epoxy 2 (Arrayit Corporation) to provide a starting matrix for DNA synthesis. Custom-designed microarrays (spot size = 17 μm) were synthesized using a photolabile deprotection strategy (Singh-Gasson et al. 1999) by Roche Nimblegen (Roche Nimblegen, Madison, WI). Reagents for synthesis (Roche Nimblegen) were delivered through the Expedite (PerSeptive Biosystems) system. For array hybridization, RNA samples (1 μg) in 1X Hybridization buffer (Roche Nimblegen) were placed in Nimblegen X12 mixer slides and incubated inside a Maui hybridization system (BioMicro^®^ Systems) for 18 hrs at 42 °C and subsequently washed according to manufacturer’s instructions (Roche Nimblegen). Arrays with fluorescently labeled RNA were imaged with a Genepix 4000B fluorescence scanner at pmt = 650 units. Secondary ion mass spectrometry analysis of microarrays hybridized with rRNA from the ^13^C- and ^15^N-labeled particle incubations was performed at LLNL with a Cameca NanoSIMS 50 (Cameca, Gennevilliers, France). A Cs+ primary ion beam was used to enhance the generation of negative secondary ions. Carbon isotopic ratios were determined on electron multipliers in pulse counting mode, measuring ^12^C^14^N- and ^12^C^15^N-simultaneously, and then ^12^C^14^N- and ^13^C^14^N-simultaneously. More details of the instrument parameters are provided elsewhere (Mayali et al. 2012). Ion images were stitched together, processed to generate isotopic ratios, and regions of interests (ROIs) of the individual probe spots extracted with the L’IMAGE software (L. Nittler, Carnegie Institution of Washington). Data were corrected for the background isotope ratios measured on probe spots hybridized with control oligonucleotides. For fluorescence, triplicate samples were combined and hybridized on the same microarray (4 total samples: FL dark, FL light, AT dark, AT light). For NanoSIMS, all replicates were analyzed separately (12 total samples).

We determined what OTUs in the different treatments were statistically isotopically enriched as described below. For each OTU and for each replicate of all treatments, ^15^N enrichment (in permil) of individual probe spots was plotted versus fluorescence and a linear regression slope, which we refer to as the hybridization-corrected enrichment (HCE), was calculated. We carried out this calculation with the ^15^N data as the ^13^C exhibits higher background and thus lower signal to noise. OTUs were considered significantly isotopically enriched if the slope minus two calculated standard errors (SE) was greater than zero and if the slope was significant based on a t-score statistic (t = slope/SE) with a p-value of less than 0.05 adjusted with the Benjamini-Hochberg false discovery rate procedure. We considered an OTU enriched in a particular treatment if two out of three replicates were statistically significantly enriched. The second type of analysis tested differential isotope labeling of the same probes with an analysis of covariance (ANCOVA), which used a standard least squares approach to determine if treatment significantly affected the slope of enrichment over fluorescence. We tested the effect of light, both on free-living and attached fractions, as well as the effect of attachment, both on light and dark incubated samples, on the HCE slope. Note that in these analyses, the model tests the contributions of three components on the isotope enrichment: 1) the experimental treatment (e.g. light vs. dark or attached vs. free-living), 2) the fluorescence, and 3) the interaction of treatment and fluorescence, so called the crossed effect. We used this latter component (the crossed effect) as an indication of the treatment having a significant effect on the isotope incorporation.

## Results

### Microbial community response to particles

Following the 72-hour incubation of Pacific Ocean microbial cells in the presence of phytodetrital particles, both cell and dissolved organic carbon (DOC) concentrations increased compared to the control (no particle) incubations (Fig. 1). Cell and DOC concentrations were positively correlated with one another for both control (r=0.95, p=0.054) and particle (r=0.83, p=0.04) incubations, suggesting that higher DOC concentrations supported increased microbial growth and vice versa. Furthermore, the dark particle incubations exhibited significantly higher cell abundances (15% increase, p=0.006) and DOC concentrations (65% increase, p=0.01) compared to the light incubations. The particles remained visibly larger and greener after incubation in the dark (Supplementary Fig. S1), suggesting light-enhanced chlorophyll degradation occurred in the light treatment.

**Fig. 1.**
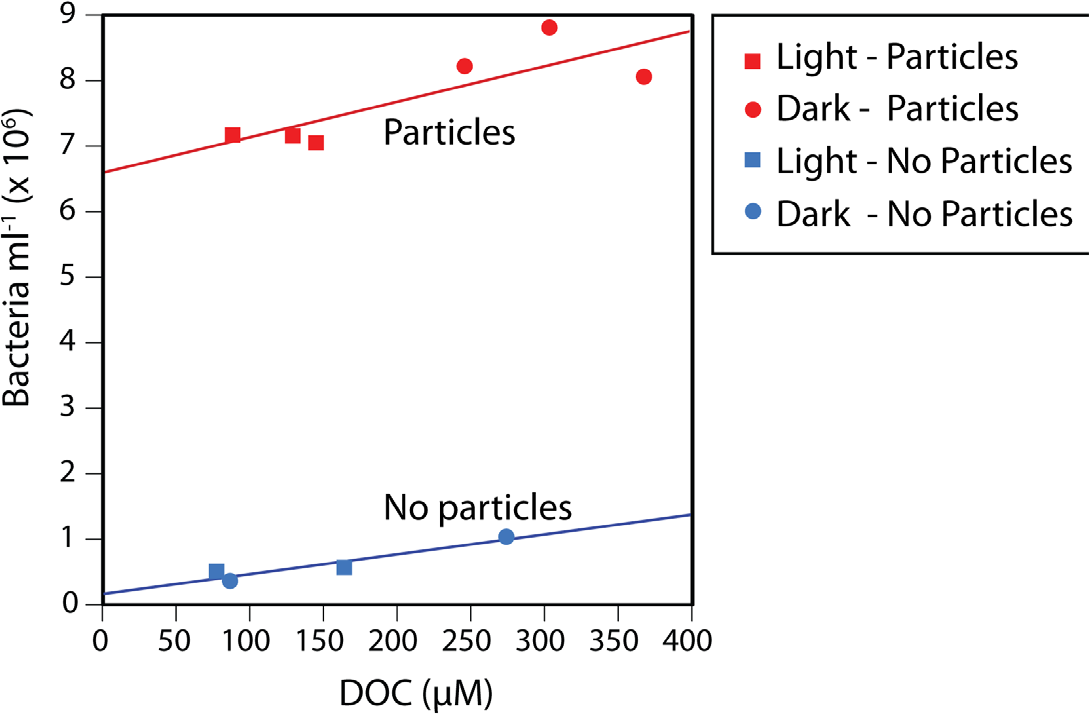
Bacterial cell abundances and dissolved organic carbon (DOC) concentrations in the control and particle addition treatments after a 72 hour incubation, showing increased DOC and bacterial concentrations in the particle incubations compared to the no-addition controls, as well as a statistically significant positive correlation between the two variables. This pattern applied to both the incubations with particles and the controls, regardless of being exposed to continuous light or darkness.

A total of 348,385 high-quality, merged 16S rRNA reads were obtained from 15 samples. After removal of chimeras (1.2% of sequences) and singletons, we obtained 1,262 OTUs clustered at 99%, including 592 that were detected in the particle-addition treatments. Microbial communities in all incubations were dominated by Gammaproteobacteria (Fig. 2), and this was more pronounced in the particle-attached fraction (>80% OTUs) compared to the free-living (~65%). The clearest patterns of differential abundance were OTUs numerically enriched in the particle treatments compared to the controls, which were numerically dominated by SAR11 OTUs (Figs. 2, S2). The particle enrichment treatments experienced a clear shift in community composition (Figs. 2, 3) with increased numbers of OTUs belonging to Flavobacteriales, Rhodobacterales, Alteromonadales, and Vibrionales (Figs. 2, S2). In general, these taxa are known copiotrophs (Buchan et al. 2014) commonly found to be associated with naturally-occurring particles in marine samples, thus we focus the rest of our analyses on the specific taxa with distinct distribution patterns within the particle addition treatments. Several differences between the free living vs. particle-attached fractions were identified (Fig. 2). Within the Gammaproteobacteria, Pseudoalteromonadaceae OTUs were in significantly higher relative abundances attached to particles compared to free-living, and also higher in the light compared to the dark. Conversely, Colwelliaceae and Vibrionales abundances increased in the free-living fractions compared to the attached fraction.

**Fig. 2.**
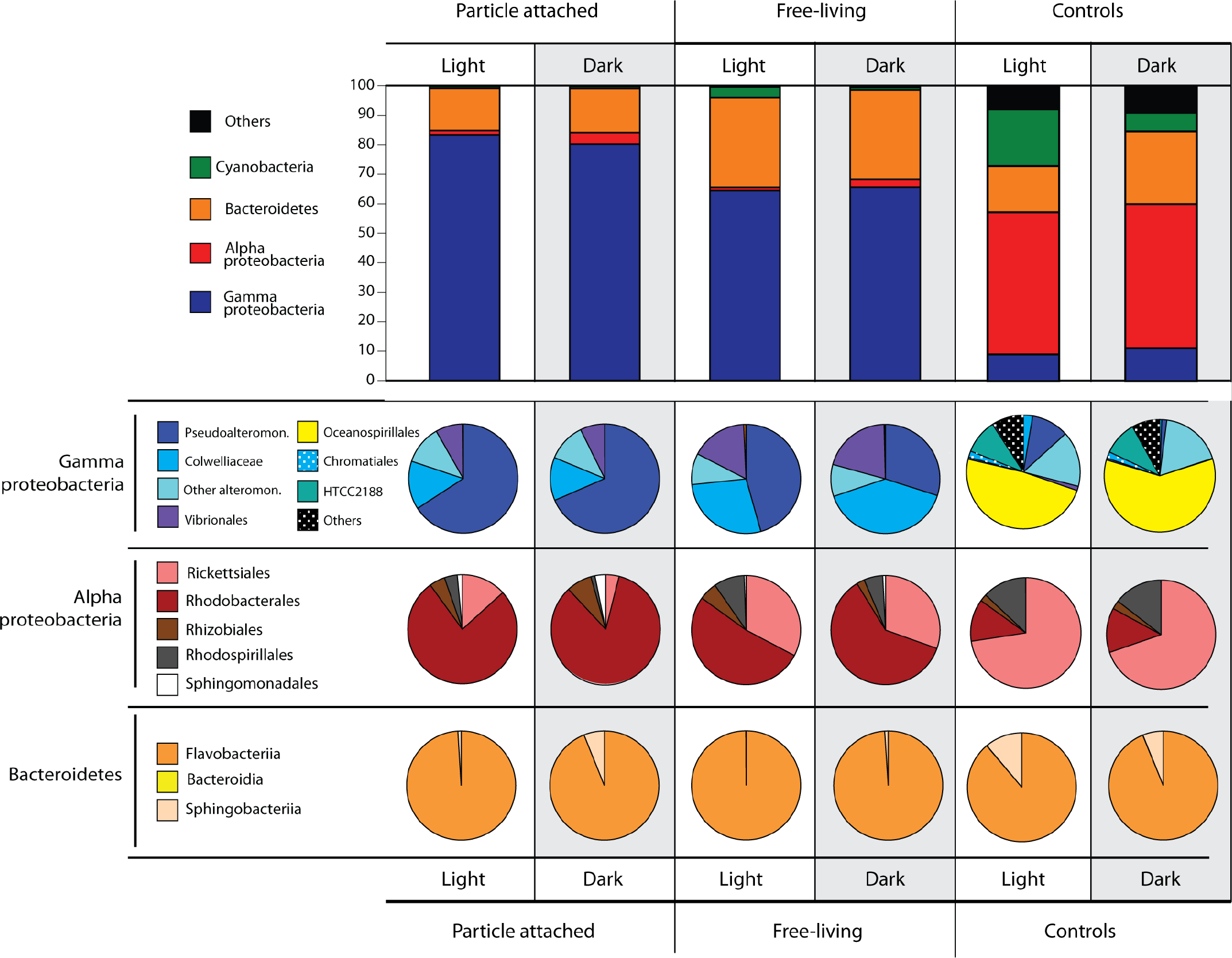
Microbial community structure identified down to the order level after 72-hour incubations in light or darkness, in the particle attached (~5 μm) and free-living (5 to 0.2 μm) fractions, and controls (no particle addition) obtained from 16S rRNA gene sequencing. Numbers refer to read percentages affiliated with the particular taxa.

Bacteroidetes were the second most abundant group detected in our samples. In contrast to many of the Gammaproteobacteria, the relative abundance of Bacteroidetes was twice as high in the free-living fractions (~30%) compared to the particle attached fractions (~15%). Within this group, >95% of the OTUs belonged to Flavobacteriia, with 5% of the reads belonging to Sphingobacteriia that were found exclusively in the dark particle-attached treatment (Fig. 2). Alphaproteobacteria and Cyanobacteria were the taxa with the overall lowest abundances in the particle addition treatments (Fig. 2). Alphaproteobacteria OTU abundances were higher in the dark treatments, Rhodobacterales being the most abundant, especially in the particle-attached treatments (>75% of all Alphaproteobacteria) compared to the free-living (50-60%). Also within Alphaproteobacteria, the Rickettsiales were most abundant in the free-living fractions of both light and dark incubations. Cyanobacteria were detected only in the light free-living samples.

The patterns described above at higher taxonomic levels were mainly driven by the appearance of individual OTUs from those taxa in the particle incubations. Many OTUs significantly responded to light regime, particle attachment, or both (adjusted p < 0.05, Fig. 3). A total of 6 OTUs were numerically enriched in the light treatments, including 3 Flavobacteria and 2 Gammaproteobacteria. We also identified 12 OTUs numerically enriched in the dark treatments, including 6 Flavobacteriales and 2 Rhodobacterales. Within the particle incubations, 32 OTUs were numerically enriched in the large particle fractions (> 5 μm), including 12 Gammaproteobacteria and 8 Flavobacteria. Twenty-one OTUs were numerically enriched in the smaller size fraction (5 to 0.22 μm), including 2 SAR11, 10 Gammaproteobacteria, and 5 Flavobacteria. Of the above mentioned OTUs, only two were significantly enriched in specific combinations of light regime and particle attachment: one Rhodobacterales (Marivita sp.) and one Pseudoalteromonas OTU, both as particle attached in the dark (Fig. 3).

**Fig. 3.**
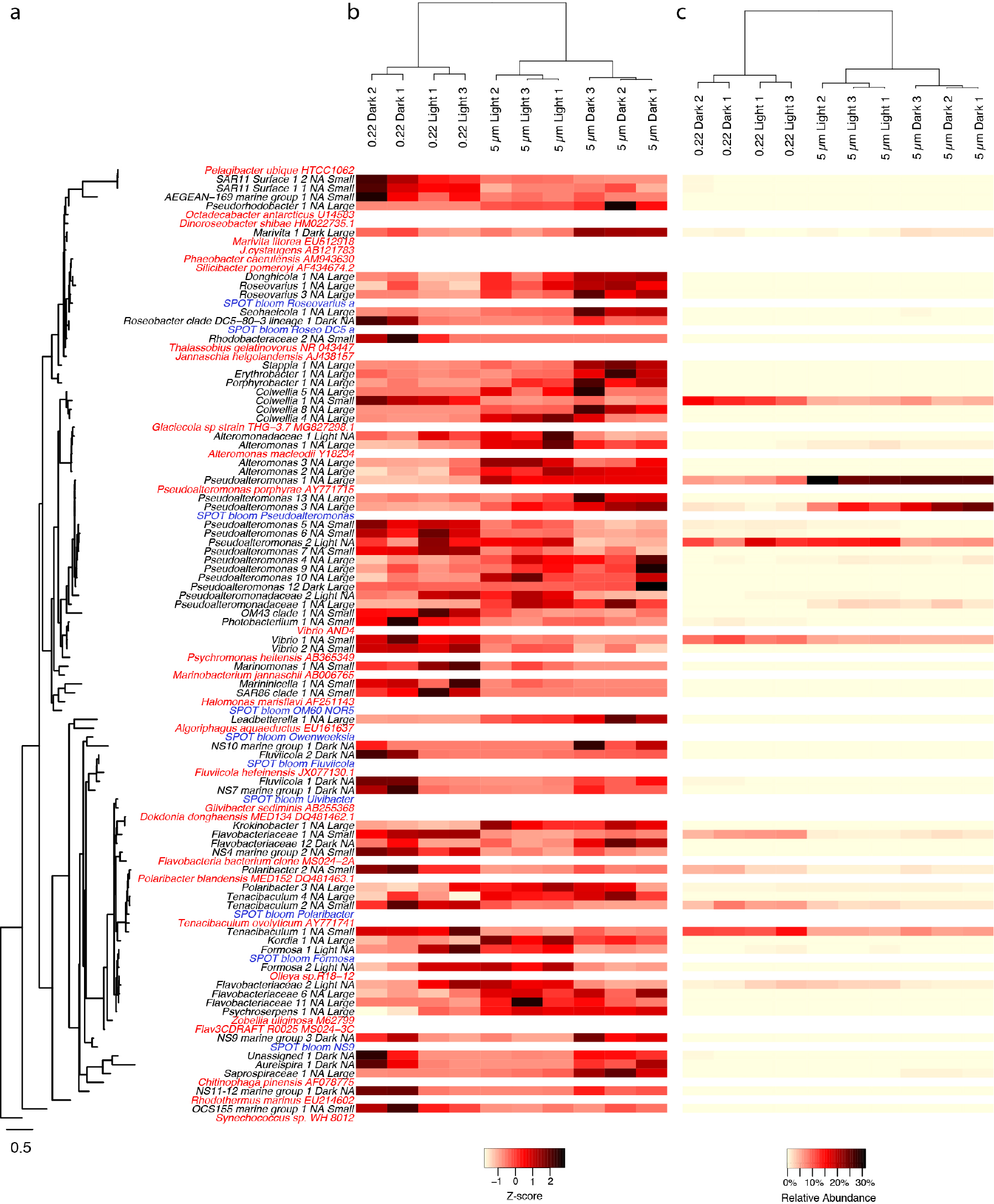
Heatmaps of OTUs that had significant shifts between the various experimental treatments. (A) OTUs are ordered by their phylogenetic relatedness of the partial 16S amplicon sequence as shown in the tree to the left, where the OTUs from the current study are in black text, reference genome sequences are in red, and OTUs from a natural phytoplankton bloom at the San Pedro Ocean Time-series location are in blue. For OTUs from the current study, they are annotated with treatments (Light or Dark, Large or Small) in which they were significant higher (p < 0.05) following the taxon name, where “Light”, and “Dark”, “Small”,“Large”, corresponds to light incubation, dark incubation, smaller size fraction (0.2 - 5 μm), or larger size fraction (>5 μm), respectively. “NA” indicates that an OTU was not significantly higher in either of the respective treatments. Relative abundances of OTUs are scaled by both their (B) z-scores and (C) relative read proportions with no transformation. Only OTUs that had relative abundances of greater than 0.25% in any of the experimental treatments are shown.

**Fig. 4.**
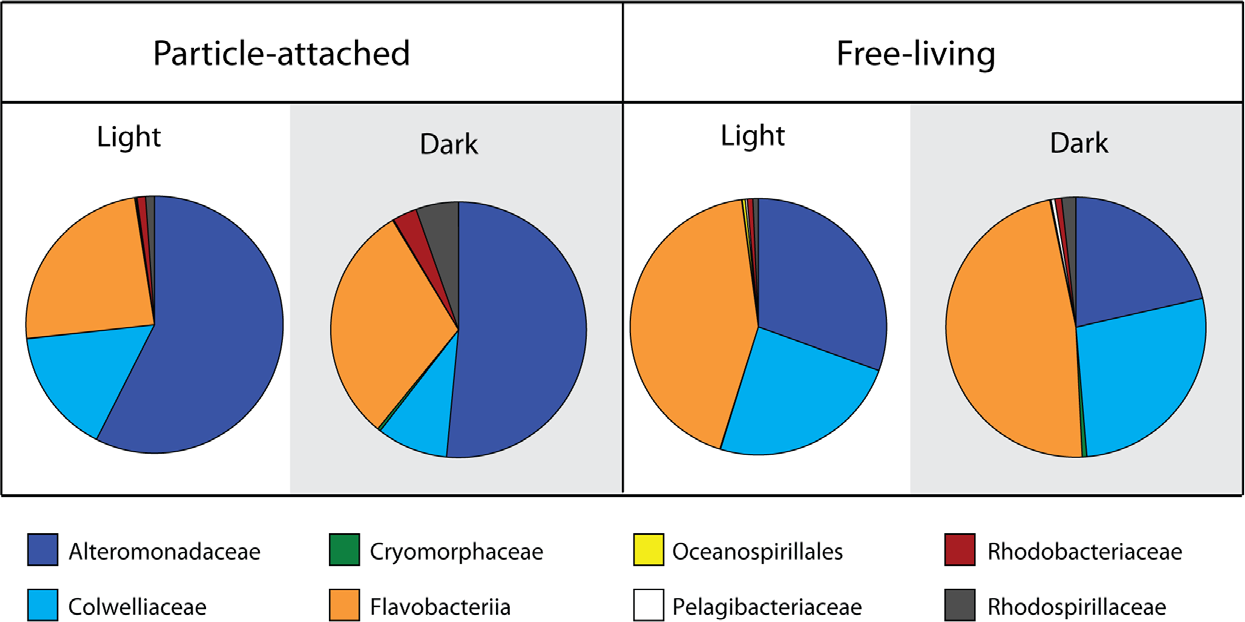
Family-level contribution of 15N incorporation from diatom-derived particles in the two treatments and size fractions. Calculations are based on taxon-specific isotope incorporation (HCE, measured by Chip-SIP) multiplied by their relative abundances based on 16S rRNA gene sequencing.

### Incorporation of isotope labeled particulate organic matter

Out of a total of 156 OTUs targeted by the microarrays, 53 OTUs were not isotopically enriched in any of the 12 samples, and another 7 were not enriched in at least two replicates of the same treatment (Fig. S3). This is a common phenomenon documented in previous Chip-SIP studies (Mayali et al. 2012) as well as density gradient SIP (Dumont and Murrell, 2005). These unenriched taxa represent organisms that were either i) in extremely low abundance in the samples or not present at all, ii) present in the sampled seawater but not growing in the bottled incubations, or iii) growing but did not incorporate C and N derived from the particles. We focus our subsequent data analyses and discussions on taxa that significantly incorporated labeled C and N from the particles. Most OTUs significantly enriched based on our criteria were, indeed, labeled in all treatments and size fractions (i.e. attached, free-living, light, dark). This shows that within the 72-hour incubation, particulate C and N was incorporated by both attached and free-living bacteria, and that this occurred both in the dark and in the light.

To obtain a taxonomic breakdown of particle-derived isotope incorporation from the different incubations and size fractions, we took the HCE values (corresponding to relative isotope incorporation) for each OTU for all replicates and averaged those values across the most abundant bacterial families. This calculation represents the relative isotope incorporation for each family, but does not take into account the abundances of those families in the samples. In order to estimate their total contribution to particulate N incorporation, we multiplied the relative isotope incorporation for each family by the percentage of reads corresponding to that family found in each of the treatments and size fractions. These calculated values should approximate the amount of isotope assimilated by the respective families in the four sample types (Fig. 4), if we assume that 16S rRNA gene read abundances correspond to relative abundances in the samples. The patterns of stable isotope incorporation in the different treatments were similar to those observed in the overall microbial community structure, retrieved by 16S iTAG sequencing. We found that the families Alteromonadaceae, Flavobacteriaceae, and Colwelliaceae dominated the isotope incorporation in the two experimental treatments with added particles. In the attached fractions, the Alteromonadaceae was particularly dominant, responsible for greater than 50% of the incorporation, both in the dark and light treatment. On the other hand, in the free-living fractions, the Flavobacteriaceae dominated isotope incorporation (48% in the dark incubations and 43% in the light; Fig. 4). As also observed by iTAG sequencing, the family Colwelliaceae incorporated the isotope mostly in the free-living fractions (24-27%) compared to the particle attached (9-16%). Other less abundant families contributed less than 1% of the isotope incorporation, with the exception of Rhodospirillaceae (5%) and Rhodobacteriaceae (3%) in the attached fraction of the dark treatment.

### Influence of light on isotope incorporation

Using covariance analysis (ANCOVA, Fig. S4), we tested which OTUs exhibited significantly distinct isotope labeling patterns according to treatment (light vs. dark) and size fraction (free-living vs. attached; Table 1). These represent taxa with significant treatment and fluorescence crossed effects on isotope labeling, with adjusted p-values <0.05. We identified 6 particle-attached OTUs that incorporated significantly more isotope in the light compared to the dark. These OTUs represented some of the most abundant taxa found in the samples according to the 16S rRNA gene sequencing data, and included the genera Pseudoalteromonas, Colwellia, and members of the Flavobacteriaceae family. One of the Flavobacteriaceae OTU (Owenweeksia) was the only OTU that also exhibited increased incorporation in the light-incubated free-living samples compared to the dark incubated samples. Seven other OTUs exhibited the opposite pattern when attached to particles, with significantly higher isotope incorporation in the dark. These included less abundant taxa, such as OTUs from the Oceanospirillales, Rhodospirillales, uncultured Flavobactericeae, uncultured Legionellaceae, and the Sva0996 marine group from the Actinobacteria. The remaining OTUs showing differential isotope incorporation were exclusively more enriched in the free-living compared to the particle attached phase (Table 1). Some of these taxa exhibited differential labeling patterns both in the light and in the dark, or in one but not the other treatment.

## Discussion

Our experimental setup was designed to use a taxon-specific, isotope labeling method (Chip-SIP) to quantify C and N incorporation by bacteria colonizing phytodetrital particles in laboratory incubations. We used freeze-thawed (dead) diatoms as model phytodetrital particles, an approach carried out in a number of previous studies (Bidle & Azam, 1999; Ploug, 2001), as they mimic natural sources of particulate C and N, which are degraded by microbial communities as they sink in the ocean. These types of studies are also valuable to understand the fate of algal detritus in outdoor raceway ponds used for biofuel production (Milledge&Heaven 2012). To our knowledge, this is the first such study to utilize isotope labeling of the algal material to trace its fate through the community. We added the impact of light in our experimental design to further examine how photoheterotrophic processes affect C and N cycling as bacteria colonized the particles. Different OTUs responded to all possible combinations of particle attachment vs free living lifestyle under light or dark conditions, at the level of 16S rRNA abundances. We further observed that some of these taxa showed differential labeling patterns, some in the light and others in the dark, as well as differential labeling between free-living and attached. The most common pattern was higher isotope incorporation in the free-living phase compared to attached (Table 1). This finding was initially rather puzzling since the isotope labeling originated from the particles and we expected isotope incorporation to be higher when particle-attached. However, previous work has shown that particle-attached bacteria exhibit greater per-cell enzyme activities but slower growth compared to their free-living counterparts, in other words, a decoupling between enzyme hydrolysis and uptake (Beier and Bertilsson 2013, Unanue et al. 1998). Furthermore, some of the taxa responsible for the first stages of particles degradation might have ended up in the small fraction after 72 hours, as we discuss below.

### Bacterial dynamics of phytodetrital particle degradation

Bacterial growth dynamics that follow algal blooms involve sequential steps of particle breakdown that are performed by different members of the microbial community. Consistent with our findings of Alteromomadaceae playing a major role in particle degradation, Sarmeto and Gasol (2012) found that Alteromonas dominated the uptake of radiolabeled diatom particles in 24-hour incubations. However, the first responders to phytodetrital particle incubations in their study were members of the Flavobacteria. Several other studies also identified Flavobacteria as the first responders after 5 hours incubations and major consumers of algal bloom biomass (Williams et al 2013, Kirchman 2002, Gomez-Pereira et al 2012). The length of our incubations (72 hours) did not allow observing the taxa that initially attached to the diatom particles during the first couple of days. Therefore, it is possible that members of Flavobacteria may have been, indeed, the first responders in our incubations, attaching to the particles and starting to degrade complex organic matter, releasing smaller breakdown products. After this first response, during which large particles are beginning to be broken down, Flavobacteria increase their relative abundances in the small fraction while the Alteromonadales dominate abundances on the particles, potentially taking advantage of less complex organic molecules produced by the Flavobacteria. We further hypothesize that this process may have been more efficient in the light due to proteorhodopsin photoheterotrophy. Flavobacteria TonB transporters, which rely on proton gradients, have been shown to be highly expressed during the peak of algal blooms (Teeling et al. 2012), suggesting a role of PR electrochemical gradients in these transport processes (Gómez-Consarnau et al. 2016). It therefore seems plausible that, in the marine environment, the first stages of diatom biomass degradation would generally be carried out mostly by photoheterotrophic Flavobacteria still in the photic zone, and that the subsequent stages of organic matter degradation would take place regardless of light as particles sink below the photic zone. Future research tracking down the transfer of biomass at different stages of an algal bloom will be necessary to better identify the role of photoheterotrophy in this first steps of algal biomass degradation.

### Light-induced microbial community response to organic particles

One of the goals of this study was to identify microbial taxa able to occupy different niches as consumers of marine organic matter in the photic zone. These include i) particle-attached phototrophs, which would take advantage of light through photoheterotrophy to enhance the transport and uptake of complex particulate organic matter, ii) particle-attached chemoheterotrophs, which include decomposers of labile complex organic matter in particles without using light, iii) free-living phototrophs, bacterial groups potentially specialized in the light-mediated utilization of smaller organic molecules that remain in seawater after being released from different sources (e.g. particle dissolution, viral lysis) and iv) free-living chemoheterotrophs, which solely rely on dissolved organic matter utilization without using light. Using our experimental incubation approach and examining relative abundance of the 16S rRNA gene and isotope incorporation differences between treatments, we were able to identify several patterns of niche preference at various taxonomic levels, including some that were only detected at the OTU level.

#### Particle-attached chemoheterotrophs vs. particle-attached photoheterotrophs

Taxa in the family Alteromonadaceae clearly dominated the phytodetrital particle incubations, with some OTUs with higher relative abundances in the attached microbial fraction and others in the free-living. Light regime, however, did not have an effect on their overall abundances (except for two OTUs with greater abundances in the light treatment). Isotope incorporation for several OTUs from this family, however, was greater in the light compared to the dark (Table 2). Alteromonadaceae are known for their opportunistic lifestyles and great capacity of growing on pulses of organic matter (Allers et al. 2007). There are virtually no phototrophs known in this family (Pinhassi et al. 2016), which would explain the general lack of growth response to light regime in our incubation experiments. Increased isotope incorporation in the light (Table 1), however, could have been a result of other microbial taxa using photoheterotrophy to start degrading high molecular weight compounds and increasing the availability of newly produced low molecular DOM for the Alteromonadaceae to subsequently incorporate. Thus, our data are consistent with the idea that photoheterotrophy led to increased C and N transfer from phototrophic to non-phototrophic bacteria. A number of other OTUs, including members of the orders Oceanospirillales, Rhodospirillales, and other Flavobacteriales, on the other hand, incorporated relatively more isotope and some were more abundant in the particle-attached dark treatment compared to the light. These taxa may be inhibited by light or the combination of light and organic matter, which can cause the formation of reactive oxygen species. Interestingly, the majority of the taxa with higher isotope incorporation in the dark correspond to sequences originally sampled from the deepest SPOT station (890 m depth; Table 1), consistent with these taxa being better adapted to dark conditions.

One surprising result from the isotope incorporation and the relative abundance data was the lack of an effect of light on the family Rhodobacteriaceae. Many of these taxa are aerobic anoxygenic phototrophs, using bacteriochlorophyll to harvest light energy (Koblizek et al. 2105). While several studies show that light seldom stimulates their growth and overall cell numbers (e.g. Ferrera et al. 2011), it can increase their substrate uptake rates (Ferrera et al. 2017). Rhodobacteriaceae are also known to follow phytoplankton blooms, taking advantage of the inputs organic matter freshly synthesized and often contained in particles (Buchan et al. 2014, Teeling et al. 2016). It is possible that due to our experimental setup consisting of 24-hour continuous light incubations, photoheterotrophic metabolism by AAP bacteria may have been inhibited in the light, as bacteriochlorophyll is synthesized during the dark cycle (Wagner-Dobler and Biebl 2006).

#### Free–living chemoheterotrophs vs. free-living photoheterotrophs

A number of studies have now shown that free-living strict chemoheterotrophy is not the most successful metabolism in surface waters, as the majority of bacteria in these environments are known free-living photoheterotrophs containing genes for proteorhodopsin phototrophy (80-100% of picoplankton genomes) or aerobic anoxygenic photosynthesis (5-30% of genomes; Dubinsky et al. 2017, Brindefalk et al. 2016, Sieradzki et al. 2018). In this study, we identified several free-living OTUs that became numerically enriched in the light compared to the dark, and one that exhibited increased relative isotope incorporation in the light versus the dark. These included several Flavobacteria OTUs, a family with numerous representatives containing PR genes (e.g. Yoshizawa et al. 2014, Gómez-Consarnau et al. 2007, Gonzalez et al. 2008, Inoue et al. 2013). However, some other Flavobacteria OTUs were enriched in the dark treatments instead, suggesting a chemoheterotrophic lifestyle by those organisms. Indeed, not all marine Flavobacteria strains contain PR genes in their genomes (Fernandez-Gomez et al. 2013), and even the ones that do seem to only respond to light when organic matter is scarce and additional scavenging capacities are needed (Gómez-Consarnau et al. 2007, 2016). A key factor in this light-induced response is that the threshold at which a population experiences substrate limitation – by type of molecule and concentration– can be different from strain to strain (Hagstrom et al. 2017). Overall, these observations within Flavobacteria reflect the high functional complexity and regulation in members of the same clade (as determined by 16S; Teeling et al. 2012). From an ecological perspective, these slight differences are likely vital to maintain separate niches in the microbial community, ultimately giving each OTU the potential to thrive in changing environmental conditions such as that experienced through an algal bloom rise and decay.

### Impact of light on organic matter utilization

Free-living bacterial cell and DOC concentrations were significantly lower in the light compared to the dark after incubation with phytodetrital particles. This suggests that either light inhibited bacterial degradation of the particles, which would result in lower bacterial growth and less dissolution of POC into DOC, or light induced more efficient particle degradation due to photoheterotrophic metabolism. If the latter, labile carbon in the light was either being i) incorporated into larger cells or ii) respired at a higher rate. Since we did not observe any obvious increases in cell size in the light, it seems more plausible that bacteria had, indeed, access to more C in the light and that more of this C was subsequently being respired. Experiments with proteorhodopsin-containing Flavobacteria strains have previously shown that light can enhance the uptake of certain organic molecules (i.e. vitamins, Gómez-Consarnau et al. 2016), which are required for growth, and therefore respiration. Under this scenario, a larger fraction of the organic matter pool would be accessible in the light due to light-enhanced DOM uptake and utilization. However, if we consider photoheterotrophy to be a dominant trait in this microbial community—as a whole— with the potential to increase bacterial fitness, why were bacterial abundances lower in the light incubations? Future research with a higher temporal resolution and performing more extensive C and N budgets will be needed to answer this question, and will require quantifying the stable isotopic signal in the particulate, dissolved and gas (i.e. CO_2_) fractions after light/dark incubations.

## Acknowledgments

Work performed by X. M. and P. K. W. was funded by the U.S. Department of Energy’s Office of Biological and Environmental Research Biofuels Science Focus Area Grant SCW1039, and work at LLNL was performed under the auspices of the US Department of Energy at the Lawrence Livermore National Laboratory under Contract DE-AC52-07NA27344. L. G. C. was funded by the Marie Curie Actions–International Outgoing Fellowships (project 253970) and the US National Science Foundation grant OCE1335269.

