## Supplementary Figures for "Influence of light on particulate organic matter utilization by attached and free-living marine bacteria"

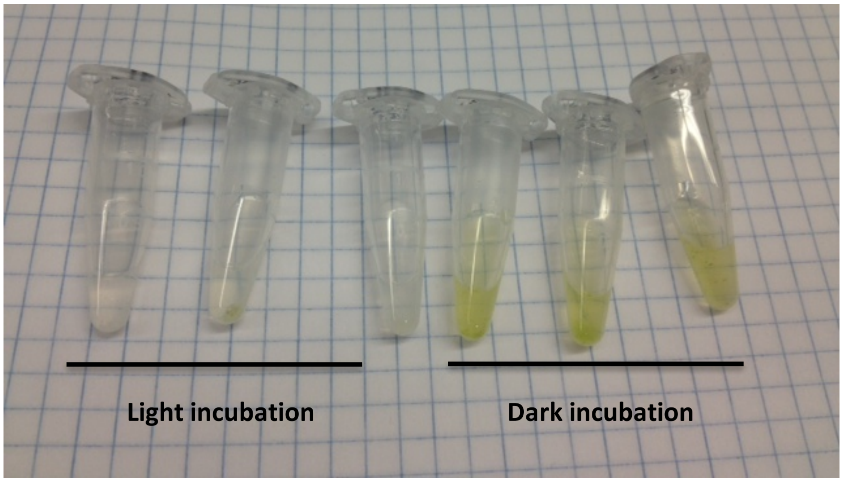


**Figure S1**. Visual appearance of the biomass in the experiment bottles after 72-hours incubation in continuous light (left) and dark (right). 5 ml samples from each incubation bottle were fixed with 10% formalin (3.7% formaldehyde), concentrated by centrifugation (10,000 rpm) and resuspended in 50 µl PBS.

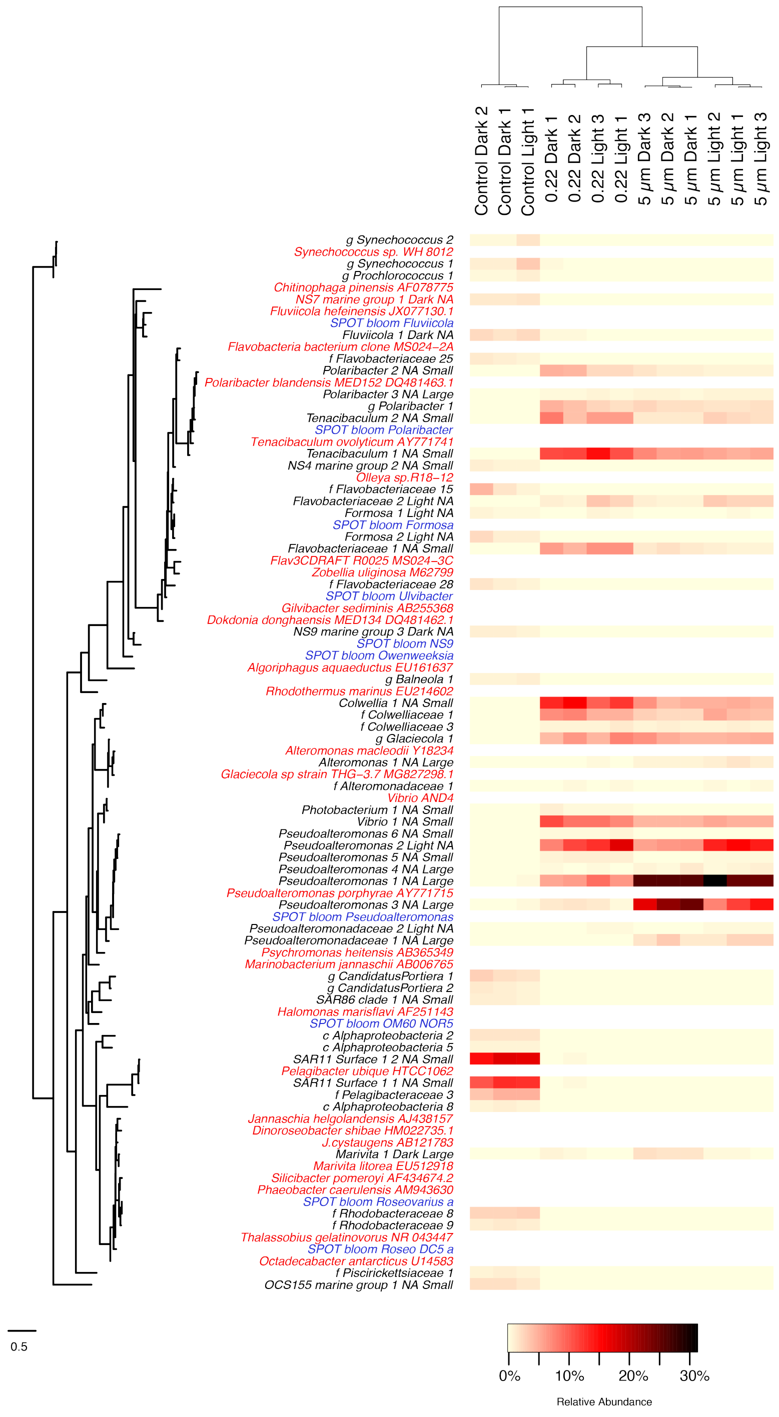


**Figure S2.** Heatmap that shows the 25 most relatively abundant OTUs, on average, in experimental treatments and controls. The two sets were mutually exclusive. Taxa identifiers in black are OTUs from the current study, in red are genomic reference sequences and in blue are OTUs from a natural phytoplankton bloom at the San Pedro Ocean Time-Series (SPOT) location.

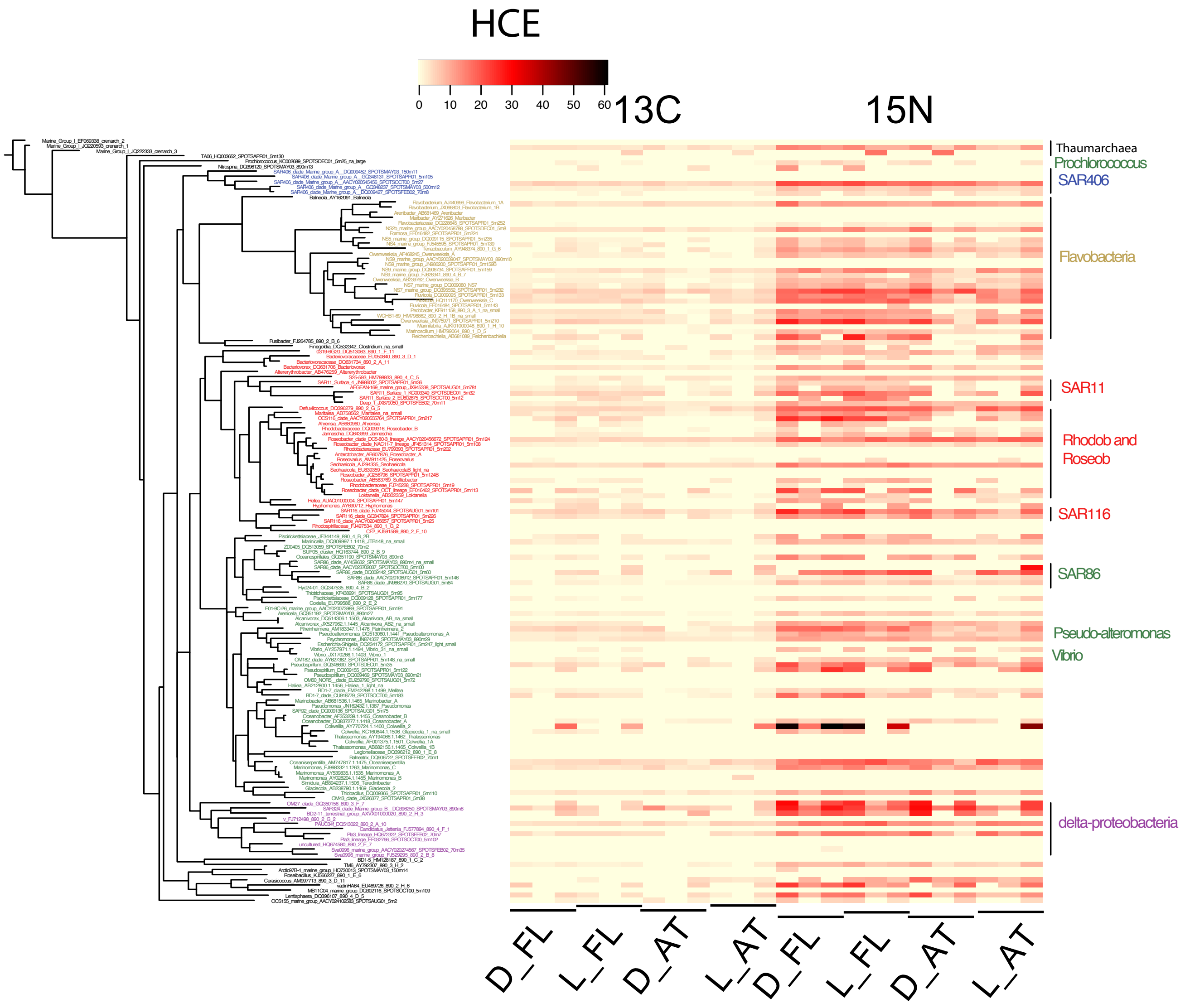


**Figure S3**. 16S rRNA gene phylogenetic tree of the 156 operational taxonomic units (OTUs) targeted by the Chip-SIP microarray. The heat map shows relative isotope incorporation calculated from both the $^{13}C$ and $^{15}N$ incorporation data from the 12 samples (D_FL = dark free-living, L_FL = light free-living, D_AT = dark attached and L_AT = light attached). OTU colors correspond to major bacterial phyla: Deferribacters in blue, Bacteroidetes in yellow, Alphaproteobacteria in red, Gammaproteobacteria in green, Deltaproteobacteria in purple.

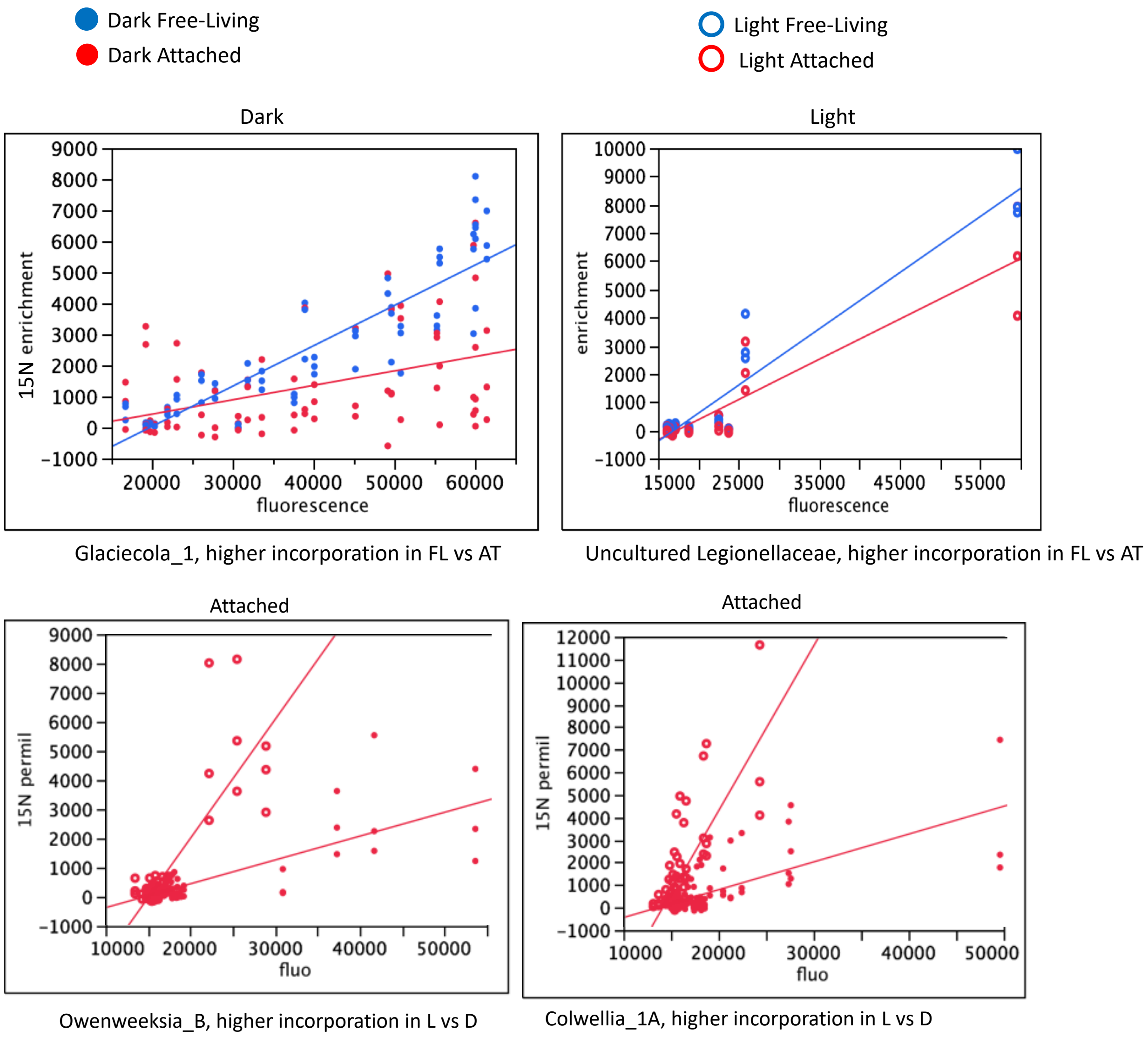


**Figure S4**. Analysis of covariance (ANCOVA) of two OTUs showing statistically significantly different treatment effects on isotope incorporation.
